# Lacer: accurate base quality score recalibration for improving variant calling from next-generation sequencing data in any organism

**DOI:** 10.1101/130732

**Authors:** Jade C.S. Chung, Swaine L. Chen

**Affiliations:** National University of Singapore, Department of Medicine, Yong Loo Lin School of Medicine, 1E Kent Ridge Road, NUHS Tower Block, Level 10, Singapore 119074; Genome Institute of Singapore, Infectious Diseases Group 3, 60 Biopolis Street, Genome, Level 6, Singapore 138672.

**Keywords:** bbase quality score recalibration, NGS, genetic variation

## Abstract

Next-generation sequencing data is accompanied by quality scores that quantify sequencing error. Inaccuracies in these quality scores propagate through all subsequent analyses; thus base quality score recalibration is a standard step in many next-generation sequencing workflows, resulting in improved variant calls. Current base quality score recalibration algorithms rely on the assumption that sequencing errors are already known; for human resequencing data, relatively complete variant databases facilitate this. However, because existing databases are still incomplete, recalibration is still inaccurate; and most organisms do not have variant databases, exacerbating inaccuracy for non-human data. To overcome these logical and practical problems, we introduce Lacer, which recalibrates base quality scores without assuming knowledge of correct and incorrect bases and without requiring knowledge of common variants. Lacer is the first logically sound, fully general, and truly accurate base recalibrator. Lacer enhances variant identification accuracy for resequencing data of human as well as other organisms (which are not accessible to current recalibrators), simultaneously improving and extending the benefits of base quality score recalibration to nearly all ongoing sequencing projects. Lacer is available at: https://github.com/swainechen/lacer.

## Introduction

Next-generation sequencing (NGS) has revealed broad insights into the prevalence and function of genetic variation, especially with respect to single nucleotide polymorphisms (SNPs) that influence human disease (Thorisson et al. 2005; Cotton et al. 2008; The 1000 Genomes Project Consortium 2012; The Cancer Genome Atlas Network 2012). Given the scale of the data, even incremental increases in accuracy have an enormous impact. Accurate SNP identification relies on accurate sequencing, which is quantified by base quality scores; unfortunately, machine-reported qualities are inaccurate (Li et al. 2009b). Therefore, recalibration of base quality scores improves downstream variant calling, mostly by excluding false positive SNPs (Li et al. 2009b; DePristo et al. 2011).

Quality score recalibration is trivial if the status of every base (correct or error) is known; the fraction of sequencing errors with a given quality score can be used to calculate the empirical (recalibrated) quality. For real sequencing data, however, erroneous bases are of course not already known. Intriguingly, all current recalibrators (Li et al. 2009b; DePristo et al. 2011; Zook et al. 2012; Cabanski et al. 2012) are strongly based on this assumption that erroneous bases are known; sequencing errors are identified as mismatches to a reference genome, excluding sites of known variants (e.g., dbSNP (Sherry et al. 2001) for humans). This assumption would be tenable if variant databases were complete, but this is also not the case (The 1000 Genomes Project Consortium 2010), and the purpose of sequencing is often to discover variants not present in existing databases. Furthermore, outside of humans and several model organisms, variant databases are not available and thus recalibration is often not done.

We have solved these logical and practical problems with a new algorithm for recalibrating base quality scores, Lacer. By comparing Lacer with GATK (DePristo et al. 2011), the current standard for base recalibration software, on four Illumina sequencing data sets (*Escherichia coli*, human, macaque and marmoset), we show that Lacer more accurately recalibrates base quality scores in the absence of complete (or any) variant knowledge, enabling application to any organism. Lacer’s more accurate recalibration in turn results in a specific increase in the confidence of true variants (based on variant quality scores), leading to more effective exclusion of false positive variants and more accurate variant calls. Finally, we demonstrate that Lacer can be applied to a wide range of NGS data sets, including different types of sequencing data (whole genome, exome, metagenome, and targeted sequencing) and data from different sequencing platforms (Illumina, Ion Torrent, and 454).

## Results

### The Lacer algorithm

Lacer (summarized in **Fig. 1A**) uses linear algebra concepts instead of assuming that correct and incorrect bases can be directly identified. Briefly, sequencing reads are mapped to a reference, and bases are binned into sets based on consensus status and depth of coverage. The sorting leverages the intuition that high coverage, consensus bases are likely to be correct, whereas singly observed, nonconsensus bases at high coverage positions are likely incorrect. Each set of bases defines a histogram of quality scores. The collection of histograms is cast as a matrix and analyzed by singular value decomposition (SVD). Assuming that correct and incorrect bases have a consistent distribution of quality scores and that the percentage of incorrect bases in each set varies, SVD will extract information that can be used to infer these distributions and the percentage of incorrect bases (for further details, see **Methods**); importantly, no individual bases are assumed to be either correct or incorrect. Finally, a Bayesian calculation based on the inferred aggregate frequency of incorrect bases and the distributions of correct and incorrect quality scores yields recalibrated quality scores.

**Figure 1:**
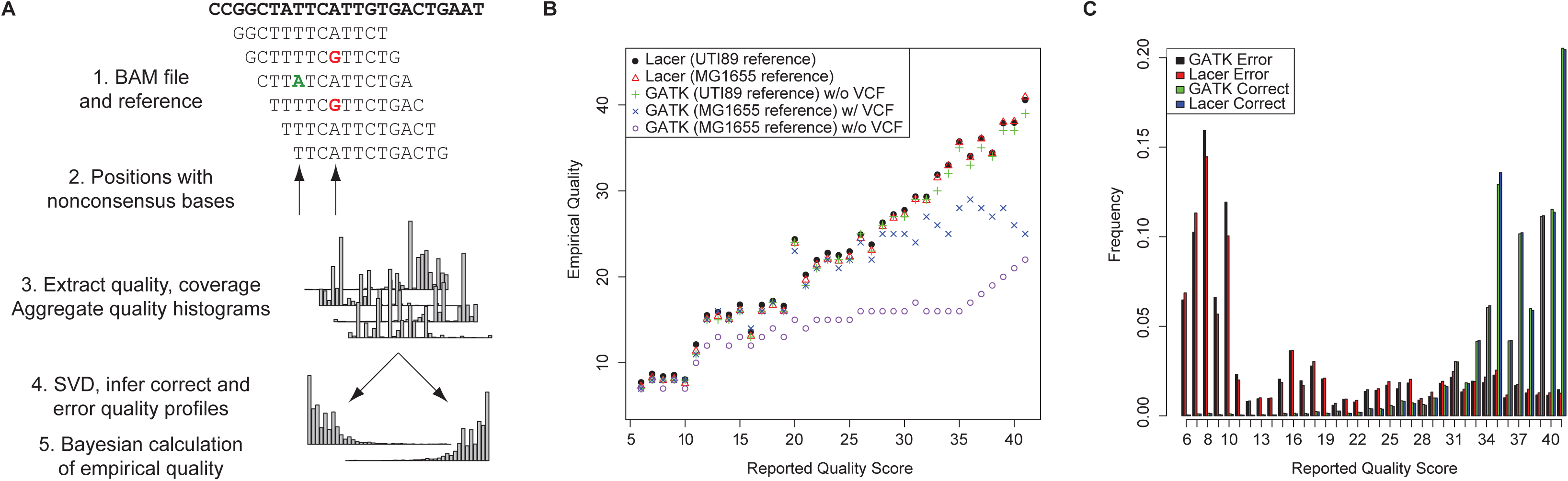
Lacer and recalibration of *E. coli* sequencing data. (**A**) Workflow of the Lacer algorithm. For details, see main text and **Methods**. (**B**) Empirical (recalibrated) quality scores plotted against reported (uncalibrated) quality scores for the UTI89 sequencing data from GATK, using UTI89 or MG1655 as a reference sequence (with or without excluding variant positions based on poorly-mapped positions and variant calls from SAMtools and LoFreq) (w/ or w/o VCF, respectively); and from Lacer, using UTI89 or MG1655 as reference and without excluding variants. Lacer recalibration using UTI89 or MG1655 as a reference matches the gold standard GATK recalibration using UTI89 as the reference. GATK recalibration using MG1655 as the reference results in generally lower recalibrated quality scores which are not fully rescued by provision of a variant database (w/ VCF). (**C**) Quality score distribution of correct and erroneous bases following recalibration of UTI89 sequencing data by GATK (with UTI89 as a reference) or Lacer (with MG1655 as a reference). The distributions generated by Lacer using the MG1655 reference match those generated by GATK using the UTI89 reference.

### Lacer matches a gold standard GATK recalibration on bacterial resequencing data regardless of reference sequence

We first compared Lacer with GATK, the current standard for base recalibration software, on an Illumina resequencing data set for *Escherichia coli* UTI89 (Chen et al. 2006), for which a complete reference sequence is available; in other words, for this data set we do know that all mismatches are indeed errors. Lacer and GATK yielded nearly identical results on UTI89 when the UTI89 genome itself was used as a reference sequence (**Fig. 1B**). Consistent with this, the quality profiles for error bases predicted by both Lacer and GATK were nearly identical (**Fig. 1C**) and matched a third method (Quake (Kelley et al. 2010)) that identifies sequencing errors using k-mer analysis (**Supplemental Fig. S1A**). GATK performs a second order recalibration incorporating covariates (sequence context and cycle number); Lacer also matched these well (**Supplemental Fig. S1B**). Therefore, in the presence of perfect information about which bases are correct and incorrect (using the known reference sequence), satisfying the assumptions made by GATK, Lacer and GATK provide concordant recalibration, which we take as gold standard recalibration for this particular UTI89 resequencing data.

In general, there is not perfect information about mutations; to mimic this, we recalibrated the same sequencing data using a different reference genome, *E. coli* MG1655 (which is ∼98% identical to UTI89). Lacer’s overall and covariate recalibration still matched the gold standard (GATK using the UTI89 reference sequence), but GATK (using the MG1655 reference sequence) reported a dramatically different recalibration, based on overall base quality recalibration, second order recalibration, and concordance of error base quality score profile with Quake (**Fig. 1B, Supplemental Fig. S1B,C**). This could not be fully mitigated by providing GATK with a maximal variant database generated by marking SNP calls from uncalibrated data as well as potentially mismapped reads (**Fig. 1B**), while Lacer’s recalibration was unaffected by exclusion of sites in this bootstrapped variant database (**Supplemental Fig. S1D**). Therefore, Lacer provides an accurate recalibration independent of an imperfect reference sequence and of knowledge of known variants, suggesting that Lacer might effectively be used to recalibrate sequencing data for any organism.

### Lacer accurately recalibrates human resequencing data without a perfect reference sequence

We therefore tested Lacer on human sequencing data for the CEPH individual NA12878 (The 1000 Genomes Project Consortium 2012), where no true gold standard recalibration is available. As seen with the UTI89 data using an imperfect reference, Lacer recalibrated the NA12878 chr1 data (mapped to the GRCh37.p13 reference genome) to higher quality scores than GATK (**Fig. 2A**) and was insensitive to the provision of known variants **(Supplemental Fig. S1E)**. To definitively verify the independence of Lacer from known variants, we restricted Lacer to only dbSNP sites and found that it produced a similar recalibration to that obtained from the full data set (**Fig. 2A**). To reconcile the differences between the Lacer and GATK recalibrations, we again used Quake as an independent method. The quality profiles for error bases predicted by Lacer and Quake were similar; however, error bases identified by GATK had consistently higher quality scores (**Supplemental Fig. S1F**), somewhat resembling the quality profile of correct bases (**Fig. 2B (inset), Supplemental Fig. S1G**).

**Figure 2:**
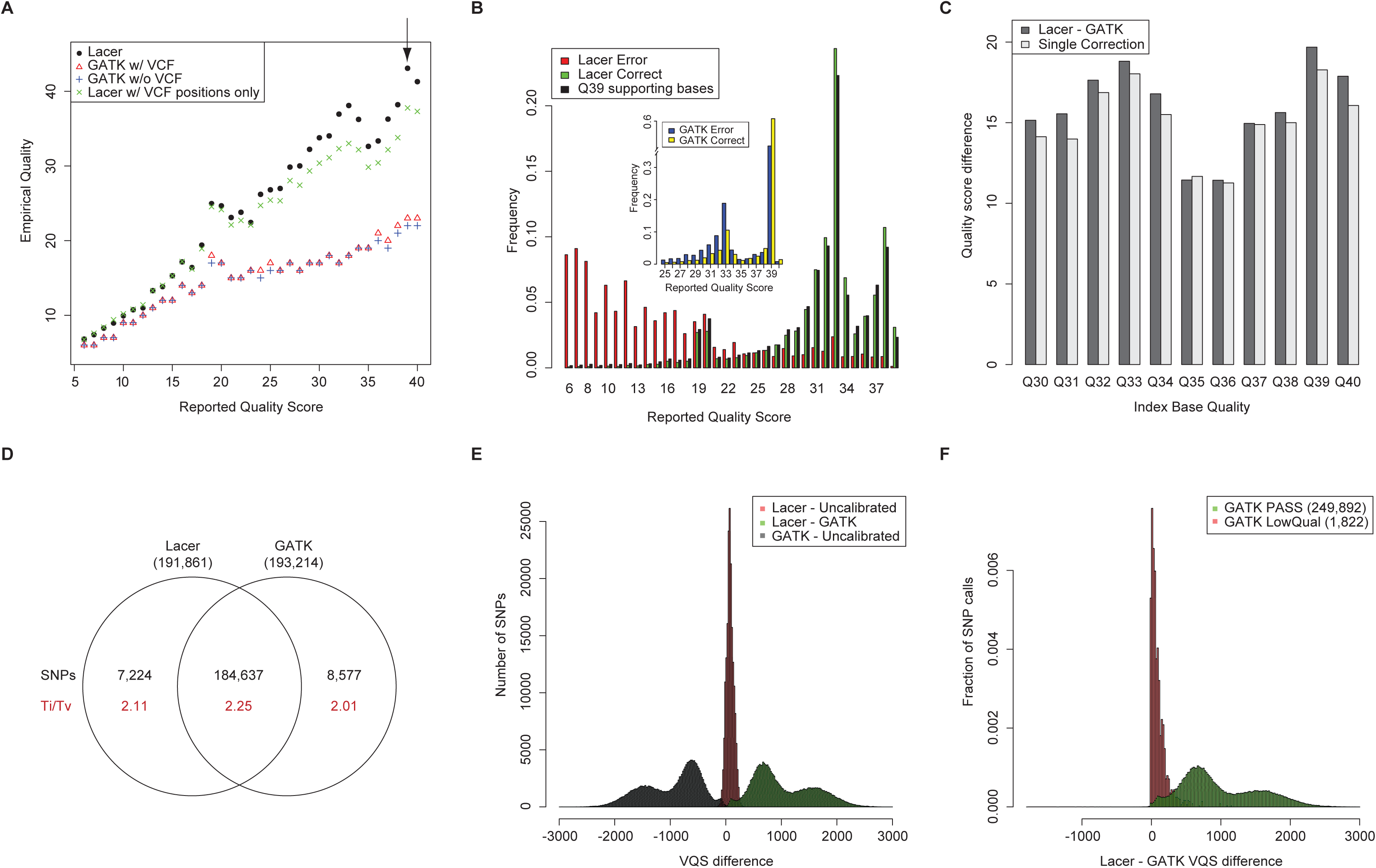
Recalibration of human sequencing data. (**A**) Recalibration of the NA12878 chr1 sequencing data using GATK, with or without excluding known variants based on dbSNP, Mills and 1000 Genomes gold standard indels, and 1000 Genomes Phase 1 indels (w/ or w/o VCF, respectively), or using Lacer, w/o VCF or with dbSNP variant positions only (VCF positions only). The Lacer recalibration results in generally higher recalibrated quality scores compared with GATK, and these are not rescued by provision of a variant database (w/ VCF). Recalibration using only VCF positions by Lacer approaches the Lacer recalibration using the full data set. (**B**) Quality score distributions of error (red) and correct (green) bases defined by Lacer recalibration and of supporting bases (black) for all non-reference, non-VCF bases with a quality score of 39 (Q39) for the NA12878 chr1 data set. All Q39 bases themselves have been excluded from this graph. The distribution for the Q39 supporting bases is similar to the distribution for the correct bases. Inset, quality score distributions for error (blue) and correct (yellow) bases defined by GATK recalibration of the NA12878 chr1 data set. The quality score distribution for error bases is enriched for high quality scores and somewhat resembles the distribution for correct bases. Only data for quality scores greater than 25 are shown. The full distribution is shown in **Supplemental Fig. S1G**. (**C**) Correction of GATK recalibration using supporting base analysis. Dark gray bars represent the difference between Lacer and GATK recalibration for each quality score. Light gray bars represent quality difference from the GATK recalibration assuming only single (unsupported), non-reference, non-VCF bases are true errors. (**D**) Venn diagram of Lacer- and GATK-recalibrated SNP calls for NA12878 chr1 identified by GATK HaplotypeCaller that passed recalibration by the GATK VariantRecalibrator (VQSR) and that also matched the high-confidence NIST call set. Total numbers of SNP calls (black) in each category and their associated Ti/Tv ratios (red) are shown. SNP calls unique to Lacer recalibration have an aggregate Ti/Tv closer to the intersection than the SNP calls unique to GATK recalibration. Comparison of VQS for SNPs identified in the uncalibrated, Lacer-recalibrated, or GATK-recalibrated NA12878 chr1 data which matched the NIST call set. Lacer recalibrated data results in overall higher VQS than that from uncalibrated (red) and GATK recalibrated (green) data on these high confidence true positive SNPs. GATK recalibrated data results in overall lower VQS than uncalibrated (gray) data on these same SNPs. (**F**) Difference in VQS between the Lacer- and GATK-recalibrated call sets for NA12878 chr1 for all high quality (PASS; green) or low quality (LowQual; red) SNPs in the GATK-recalibrated call set. Lacer recalibrated data results in a mild increase in VQS on low quality SNPs (as called by GATK recalibrated data), but this is small compared to the increase in VQS for high quality SNPs.

We suspected that the discrepancy between the GATK and Lacer recalibrations was because GATK misclassified some correct bases; an incomplete variant database would result in bases supporting novel variants being classified as incorrect, and in the extreme, without a variant database, GATK indeed recalibrates to lower quality (**Fig. 2A**). We therefore examined bases of a given quality score (for example, Q39 (arrow in **Fig. 2A**)) that did not match the reference and that were not in the variant database. These are bases that would be called errors by GATK. We extracted bases that supported these “errors” – bases from different reads but mapping to the same position with the same nucleotide. The quality score distribution of these supporting bases was similar to the quality score distribution of correct bases as predicted by Lacer (**Fig. 2B**), suggesting that these bases were largely correct. However, a fraction of the Q39 “errors” was not supported (“single”). This fraction was relatively small for both the NA12878 chr1 data set and for UTI89 mapped to the MG1655 genome, but was ∼90% for UTI89 mapped to the UTI89 genome (**Supplemental Fig. S2A**). The increase in single bases was not due to generally lower coverage at these positions (**Supplemental Fig. S2B**). By hypothesizing that single bases were the only true incorrect bases, we could derive the difference in recalibrated quality scores between Lacer and GATK (**Fig. 2C, Supplemental Fig. S2C**). In other words, amending error bases identified by GATK to include only single bases and to exclude all supported bases (whose overall quality profile was similar to that for correct bases) resulted in concordant recalibration with Lacer. Given the results using both *E. coli* and human data, we conclude that Lacer provides a more robust and more accurate recalibration regardless of organism.

### Lacer recalibration improves SNP calling on human sequencing data

Using the GATK best practices pipeline (DePristo et al. 2011; Van der Auwera et al. 2013) to call SNPs on NA12878 chr1, we next compared no base quality recalibration to Lacer and GATK (**Table 1**). Recalibration with either Lacer or GATK resulted in ∼250,000 final calls; without calibration, an additional 45-57,000 SNPs were identified. SNPs excluded by Lacer appeared to be of lower quality, based on the transition-transversion ratio (Ti/Tv) of 1.49 for Lacer compared to 1.66 for GATK. The expected value for true positives is ∼2. On a high confidence, true positive SNP call set (NIST (Zook et al. 2014)), Lacer and GATK both predicted ∼7,000-8,000 unique SNPs each; the Ti/Tv for the unique Lacer SNPs was closer to the value in the intersection and higher than the value for the unique GATK SNPs (**Fig. 2D**). Therefore, recalibration with Lacer provides more effective exclusion of false positive SNPs (the primary benefit of recalibration) than with GATK.

**Table 1:** Impact of base quality score recalibration by Lacer and GATK on SNP calls from the NA12878 chr1 (GATK HaplotypeCaller; filtered by VQSR), Tibetan macaque chr1 (LoFreq), and common marmoset exome (LoFreq) data sets. VQSR: variant quality score recalibration; recal: recalibration; LRA: local realignment.

| Call Set | NA12878 chr1 |  |  |  | Tibetan macaque chr1 |  |  |  | Common marmoset exome |  |  |  |
| --- | --- | --- | --- | --- | --- | --- | --- | --- | --- | --- | --- | --- |
|  | No. of SNPs |  | Ti/Tv |  | No. of SNPs |  | Ti/Tv |  | No. of SNPs |  | Ti/Tv |  |
|  | Lacer | GATK | Lacer | GATK | Lacer | GATK | Lacer | GATK | Lacer | GATK | Lacer | GATK |
| Raw reads, all calls | 305,220 | 305,220 | 2.00 | 2.00 | 1,463,349 | 1,463,349 | 1.62 | 1.62 | 7,089,824 | 7,089,824 | 2.06 | 2.06 |
| Unique to raw read calls | 2,902 | 12,859 | 0.70 | 0.86 | 29,771 | 34,390 | 0.57 | 0.66 | 20,276 | 69,222 | 1.05 | 1.36 |
| Unique to +recal/+LRA | 2,187 | 3,173 | 1.01 | 1.34 | 40,328 | 66,202 | 0.68 | 0.65 | 62,459 | 229,667 | 1.17 | 1.17 |
| +recal/+LRA, all calls | 304,505 | 295,534 | 2.01 | 2.07 | 1,473,906 | 1,495,161 | 1.62 | 1.59 | 7,132,007 | 7,250,269 | 2.05 | 2.03 |
| Filtered | 57,303 | 44,853 | 1.49 | 1.66 | 204,625 | 212,664 | 0.60 | 0.58 | 4,763,917 | 4,867,640 | 1.95 | 1.92 |
| Final call set | 247,202 | 250,681 | 2.16 | 2.16 | 1,269,281 | 1,282,497 | 1.91 | 1.89 | 2,368,090 | 2,382,629 | 2.27 | 2.26 |
| Unique to final call set | 12,030 | 15,509 | 1.84 | 1.82 | 7,080 | 20,296 | 1.70 | 0.94 | 39,237 | 53,776 | 1.86 | 1.59 |

To discover why Lacer excluded more false positive SNPs, we examined the final variant quality scores (VQS) of SNPs following variant quality score recalibration (VQSR). Lacer recalibration produced overall higher VQS than GATK or uncalibrated SNP calls on the high-confidence NIST SNPs (**Fig. 2E**). Notably, GATK resulted in generally lower VQS than uncalibrated data. To verify that Lacer was not simply elevating the quality scores of all bases, we took the set of final high and low quality SNP calls from GATK recalibration and examined their VQS in the GATK, Lacer, and uncalibrated data. Lacer had increased the VQS on high quality SNPs without affecting VQS for low quality SNPs, as expected, when compared with GATK or no recalibration. (**Fig. 2F, Supplemental Fig. S2D**). Again, GATK, in contrast, reduced the VQS specifically on high quality SNPs compared with uncalibrated data, the opposite of what would be expected (**Supplemental Fig. S2E**). We noticed a bimodal distribution in the VQS differences between GATK and uncalibrated or Lacer recalibrated data (potentially due to bimodal base quality scores after GATK recalibration) (**Supplemental Fig. S2F**). We therefore performed the entire analysis on another chromosome (chr19) that didn’t have this artifact and saw similar improved results for Lacer (**Supplemental Fig. S3, Supplemental Table S1**). Thus, Lacer recalibration results in a specific increase in the confidence of true SNPs without inflating the confidence of false SNPs, as would be expected from accurate recalibration, while GATK has the opposite effect.

### Lacer recalibration of non-human sequencing data results in improved SNP calls

To further show that Lacer can effectively recalibrate the sequencing data for any organism in the absence of a perfect reference, we tested Lacer on two non-human primate data sets, Tibetan macaque (*Macaca thibetana*) (Fan et al. 2014) and common marmoset (*Callithrix jacchus*). Lacer recalibrated the Tibetan macaque chr1 data (mapped to the rhesus macaque (*Macaca mulatta*) reference) to higher quality scores than GATK; given that known variants in this organism are likely incomplete, this was similar to the result obtained with the human data and the *E. coli* data mapped to an imperfect reference (**Fig. 3A, compare with Fig. 1B and Fig. 2A**). Furthermore, error bases identified by GATK again had consistently higher quality scores, and the quality profile resembled that of correct bases predicted by both GATK and Lacer (**Supplemental Fig. S4A**,**B**). In the presence of a complete reference sequence (calJac3 (The Marmoset Genome Sequencing and Analysis Consortium. 2014)), Lacer and GATK again yielded concordant results on the common marmoset exome data (**Fig. 3B**). To mimic the absence of perfect information about mutations, we recalibrated the same sequencing data using the rheMac3 reference genome. Again Lacer’s recalibration matched the recalibration on the perfect data, but GATK’s recalibration was very different (**Supplemental Fig. S4C,D**). Unfortunately, we could not independently verify the quality score profiles of error bases predicted by Lacer using Quake due to the low coverage (<20×) of these data sets.

**Figure 3:**
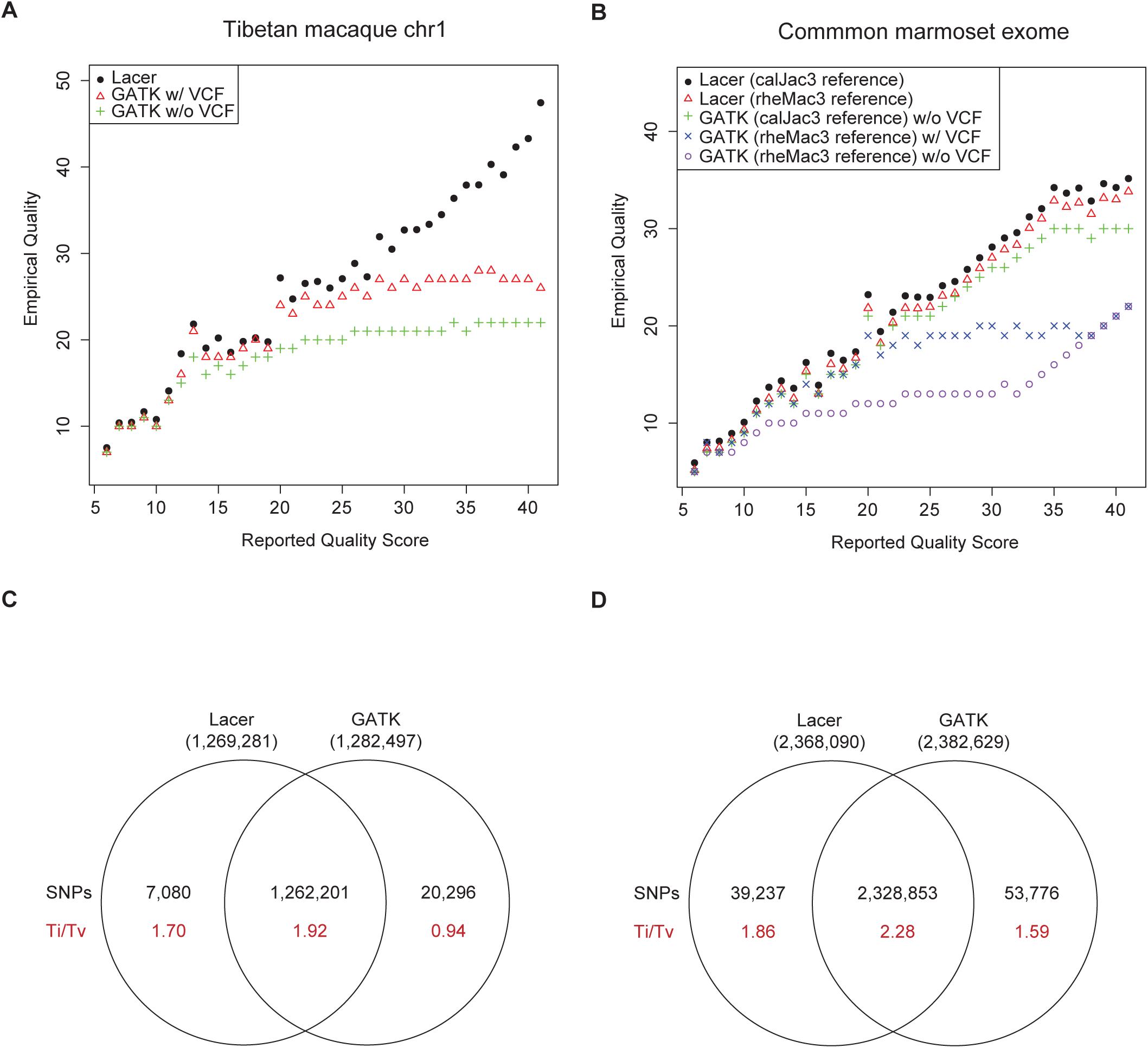
Recalibration of non-human primate sequencing data. Recalibration of (**A**) Tibetan macaque chr1 sequencing data using Lacer or GATK, with or without excluding known variants based on SAMtools and LoFreq (w/ or w/o VCF, respectively). GATK recalibration produces lower quality scores that are only partially increased by exclusion of known variants, and (**B**) common marmoset exome sequencing data using Lacer or GATK, with or without excluding known variants based on SAMtools and LoFreq (w/ or w/o VCF, respectively), and the calJac3 or rheMac3 reference genomes. Lacer recalibration using calJac3 or rheMac3 as a reference matches the gold standard GATK recalibration using calJac3 as the reference. GATK recalibration using rheMac3 as the reference results in generally lower recalibrated quality scores which are not fully rescued by provision of a variant database. Venn diagram of Lacer- and GATK-recalibrated final SNP calls for (**C**) Tibetan macaque chr1 and (**D**) common marmoset exome data, as identified by LoFreq. Total numbers of SNP calls (black) in each category and their associated Ti/Tv ratios (red) are shown. SNP calls unique to Lacer recalibration have an aggregate Ti/Tv closer to the intersection than the SNP calls unique to GATK recalibration for both data sets.

We next used the GATK best practices pipeline and the quality score-aware SNP caller, LoFreq (Wilm et al. 2012), to call SNPs on these data sets (**Table 1**). For the Tibetan macaque chr1 data, recalibration with either Lacer or GATK resulted in ∼1,300,000 final calls; without calibration, an additional 180-190,000 SNPs were identified. Recalibration by Lacer led to a slightly higher Ti/Tv in the higher call set than GATK; but the SNPs unique to Lacer (∼7,000) had a much higher Ti/Tv (1.70) than SNPs unique to GATK (∼20,000, Ti/Tv 0.94) (**Fig, 3C**). For the common marmoset exome data, recalibration with either Lacer or GATK resulted in ∼2,400,000 final calls; without calibration, an additional ∼4.6 million SNPs were identified. Once again, SNPs unique to Lacer (∼39,000) had a higher Ti/Tv ratio (1.86) than SNPs unique to GATK (∼54,000; 1.59) (**Fig. 3D**). Therefore, accurate recalibration by Lacer provides substantial improvement to SNP calling without requiring a variant database or a perfect reference sequence.

## Discussion

Lacer solves the two primary practical and logical problems with current recalibrators and is thus the first and only way at present to correctly recalibrate most NGS data (including exome, metagenome, targeted sequencing, and data across multiple sequencing platforms) (**Supplemental Fig. S5**). Lacer (i) does not require a variant database, (ii) can tolerate an imperfect reference sequence (Lacer may in future be extended to utilize k-mer analysis, potentially eliminating the need for reference sequences entirely), and (iii) is more accurate across a wider range of data sets (including human) than GATK, resulting in better downstream variant calls. Lacer is the first algorithm to extend the increased accuracy of variant calling from base quality recalibration to all non-human resequencing projects.

Current base recalibrators assume that any mismatch to a reference genome is a sequencing machine error. In reality, these mismatches may also include true variants, PCR-amplification errors, and data processing errors (e.g. alignment errors). These three classes of mismatches arise from sources outside of the sequencing process itself; in other words, if there is a true unknown variant, current recalibrators will treat those bases as errors despite the sequencing machine having sequenced them correctly. A complete variant database will only prevent misidentification true variants as error bases, and mapping scores could be used to minimize the impact of alignment errors. However, errors introduced during PCR cannot be eliminated by current recalibrators. In practice, we find that current recalibration algorithms significantly (10- to 100-fold) underestimate empirical quality scores, especially among high quality bases, even when provided with a variant database. Importantly, these high quality bases account for the majority of the data (see, for example, the black bars in **Fig. S2F**). In contrast, the Lacer algorithm directly extracts quality scores and aggregate error probabilities based on the assumption that correct and incorrect bases have different quality score profiles. The calculation of consensus bases effectively eliminates unknown true variants as a source of error and seems to mitigate against mapping errors. Based on the bacterial and human sequencing data, single (unsupported), non-consensus bases are the majority of true sequencing errors; in other words, the majority of supported bases, even with only 2 supporting bases at high coverage (>50×) positions, are actually correct. These bases generally do not result in a SNP call by current algorithms, and we therefore hypothesize that these bases may be due to PCR amplification or other unknown sources of error. Regardless, Lacer is robust to all of these potential sources of error that affect current recalibrators, and in so doing is the first recalibrator to effectively isolate and correct the errors introduced during the sequencing process itself, which is precisely what the base quality scores should be measuring.

Lacer specifically increases the confidence (or VQS) of high quality SNPs without affecting the confidence of low quality SNPs. Intriguingly, GATK’s recalibration resulted in a *reduction* in VQS for high quality SNPs; again, incomplete variant knowledge leads to the misclassification of error bases, a reduction in the empirical quality score, and the concomitant decrease in VQS. Interestingly, these lowered VQS did not significantly impact GATK’s ultimate SNP calls from the human data, based on Ti/Tv ratios and comparison to the high confidence NIST SNP set. This may be a result of the fact that GATK’s best practices pipeline is tuned for the identification of genetic variants in human sequencing data; the availability of large human training data sets enables more effective prediction of SNPs. The tuning of GATK for human data, however, limits its utility for non-human sequencing. Indeed, when applied to non-human data sets, Lacer’s consistent performance relative to GATK is apparent; not only are false positive SNPs more effectively excluded, but unique SNPs predicted from Lacer-recalibrated data also have quality metrics more consistent with that of true positive SNPs. Use of the LoFreq SNP caller (instead of the GATK HaplotypeCaller) for these data sets more effectively captures the benefit of accurate recalibration, since LoFreq takes into account base quality scores for SNP prediction. Furthermore, VQSR is not applicable to non-human data sets due to the lack of training data, which substantially reduces the accuracy of SNP identification.

Currently, there are very few methods to objectively and computationally assess the accuracy of variant calls; since Lacer uses the distribution of quality scores to calculate aggregate numbers of correct and incorrect bases, it could conceivably accomplish this when combined with the analysis of supporting bases. A further extension could enable the systematic assessment of variant calling covariates to identify rigorous filtering criteria. Finally, future SNP callers may gain additional power and accuracy by directly incorporating these insights about quality scores.

## Methods

### Lacer algorithm – theory

We first assume that correct and incorrect bases have consistent but different reported quality distributions. The quality score distribution of correct bases is denoted as *C* = { *c*_*j*_ } and that of incorrect bases as *E* = { *e*_*j*_ }. *C* and *E* are probability distributions over the set of assigned quality scores and therefore satisfy ∑_*j*_ *c*_*j*_= ∑_*j*_ *e*_*j*_= 1. Considering a set *s* containing *n* bases, *s* will have a fraction *p* correct bases and (1 - *p*) incorrect bases. Since correct and incorrect bases have consistent distributions, *s* is sampled from the quality score distribution:

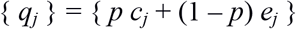

also satisfying ∑_*j*_ *q*_*j*_ = 1. Clearly, as *n* increases, the sampling error of { *q*_*j*_ } decreases. Given a collection of sets *S* = { *s*_*i*_ }, each with *n* bases and containing a fraction *p*_*i*_ of correct bases, a collection of quality profiles *Q* = { *q*_*ij*_ } can be created and represented as a matrix.

Crucially, for any given *i*:

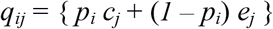

Considering this as a vector, and given *c*_*j*_ and *e*_*j*_, differences between *q*_*ij*_ can be parameterized with one variable, *p*_*i*_. Although *p*_*i*_, *c*_*j*_ and *e*_*j*_ are *a priori* unknown, providing *p*_*i*_ are not all identical, a singular value decomposition (SVD) can extract the covariance between the individual quality score variations and enable deduction of *p*_*i*_. Explicitly:

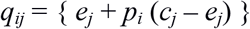

An SVD produces *c*_*j*_ – *e*_*j*_ (defined up to a sign) as the first eigenvector, while *p*_*i*_ is related (up to addition of a constant due to data centering) to the eigencoordinates in the first eigendimension. Careful construction of the sets of bases *S* = { *s*_*i*_ } provides a reasonable assurance that *p*_*i*_ spans close to the entire range from 0 to 1; this is done as described in the main text by sorting bases by consensus status (including number of supporting bases) and depth of coverage at that position. Thus, *c*_*j*_ and *e*_*j*_ can be directly calculated from the SVD results as min(*x*_*1i*_) *v*_*1j*_ and max(*x*_*1i*_) *v*_*1j*_ (adjusted for centering), respectively, where *x*_*1i*_ are the coordinates in the first SVD dimension and *v*_*1j*_ are the components of the first SVD eigenvector.

Once *c*_*j*_ and *e*_*j*_ are known, a Bayesian calculation determines the relationship between reported and empirical quality. Empirical quality can be recorded as:

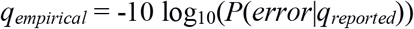

By Bayes theorem:

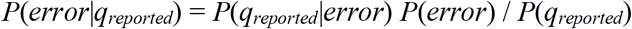

Tabulation of the overall distribution of quality scores in the sequencing data set gives *P*(*q*_*reported*_). *P*(*q*_*reported*_|*error*) is given by *E* = { *e*_*j*_ }. Importantly, since each set *s*_*i*_ contains *n* bases and the SVD provides *p*_*i*_, the total number of error bases can be calculated as ∑_*i*_ *n* (1 – *p*_*i*_) (without classifying individual bases as correct or incorrect). Division by the total number of bases then yields *P(error)*.

### Sequencing data and data processing

#### Human data sets

The sequencing data set for the CEPH individual NA12878 (sequenced on an Illumina Genome Analyzer II and aligned to the GRCh37.p13 human reference genome) was publically available from the 1000 Genomes database (The 1000 Genomes Project Consortium 2012). The high-confidence variant call set used for benchmarking was available at the NCBI Genome in a Bottle ftp site (Zook et al. 2014): NISTIntegratedCalls_14datasets_131103_allcall_UGHapMerge_HetHomVarPASS_VQSRv 2.18_all_nouncert_excludesimplerep_excludesegdups_excludedecoy_excludeRepSeqSTRs_n oCNVs.vcf. The BAM file for the NA12878 exome data set (sequenced on the Illumina Genome Analyzer IIx and aligned to the NCBI36 human reference genome using Maq) was available from the 1000 Genomes Project database (The 1000 Genomes Project Consortium 2012). The BAM file for the TruSeq Amplicon Cancer Panel targeted sequencing data (sequenced on an Illumina MiSeq and aligned to the NCBI36 human reference genome) was available from the BaseSpace Public Data repository (https://basespace.illumina.com/run/358358).

Sequencing data were aligned to the corresponding reference genomes using BWA (Li and Durbin 2009). SAMtools (Li et al. 2009a) was utilized for alignment, sorting, and indexing of sequencing reads. The aligned data was then processed following the GATK (version 2.8-1) best practices workflow (DePristo et al. 2011; Van der Auwera et al. 2013). For the NA12878 whole genome sequencing data set, a variant database for GATK’s recalibration was generated by including variant positions from dbSNP (BUILD_ID 137, reference GRCh37.p10), Mills and 1000 Genomes gold standard indels (reference GRCh37.p10), and 1000 Genomes Phase 1 indels (reference GRCh37.p10), which were downloaded from the GATK resource bundle (version 2.5). For the NA12878 exome and TruSeq Amplicon Cancer Panel targeted sequencing data sets, a variant database for GATK’s recalibration was generated using the GATK resource bundle for the NCBI36 reference genome. Blank VCF files for all data sets were generated by including mock VCF headers and fields alone.

#### Non-human data sets

For the UTI89 Illumina sequencing data set, UTI89 was grown in LB broth overnight at 37°C with agitation. Genomic DNA was purified using the Wizard genomic DNA purification kit (Promega). DNA was sheared using the Covaris S2x and size selected on a Pippin Prep (Sage Science) to isolate 400-500 bp fragments. Illumina sequencing libraries were constructed using the Illumina TruSeq kit using manufacturers protocols, then sequenced on a HiSeq 2000 with 2×76 paired end reads. For the UTI89 Ion Torrent sequencing data set, UTI89 was grown in LB broth overnight at 37°C with agitation. Genomic DNA was purified using the Wizard genomic DNA purification kit (Promega). Ion Torrent sequencing libraries were constructed using the Ion Xpress Plus Fragment Library kit (Life Technologies) according to the manufacturer’s protocols, then sequenced on an Ion 316 chip. The MG1655 454 data set (sequenced on a Roche 454 GS FLX Titanium System) was available from the SRA database (SRX255226; www.ncbi.hlm.nih.gov/sra). The UTI89 and MG1655 sequencing data were aligned to the *E. coli* UTI89 (NC_007946) or *E. coli* MG1655 (NC_000913) reference genomes. The SRS017191 human metagenome (sequenced on an Illumina Genome Analyzer IIx and aligned to the *Bacteriodes vulgatus* ATCC 8482 (NC_009614) reference genome) data set was available from the Human Microbiome Project database (The NIH HMP Working Group 2009). The Tibetan macaque (*Macaca thibetana*) data set (sequenced on an Illumina HiSeq 2000) was available from the SRA database (SRX373102); sequences were trimmed to equal lengths and aligned to the *Macaca mulatta* (rheMac3; rhesus macaque) reference genome. The common marmoset (*Callithrix jacchus*) exome data set (sequenced on an Illumina HiSeq 2000 and aligned to the calJac3 (*Callithrix jacchus*; The Marmoset Genome Sequencing and Analysis Consortium. 2014) or rheMac3 reference genomes) was available from the SRA database (SRX375642).

The sequencing data were aligned to the corresponding reference genomes using BWA. SAMtools was utilized for alignment, sorting, and indexing of sequencing reads. With the exception of the Tibetan macaque and marmoset data sets, which were processed following the GATK best practices workflow, all aligned data were recalibrated directly. A variant database for GATK’s recalibration was generated by including variants called by SAMtools, LoFreq (Wilm et al. 2012), and where indicated, positions showing poor mappability.

### Supporting base analysis

The SAMtools Perl API and custom Perl scripts were used to perform pileups and extract nonconsensus bases, their corresponding quality, and the same data from supporting bases (identical bases from different reads) at positions outside of dbSNP from the aligned NA12878 data. Using custom R scripts, histograms of supporting base qualities were plotted and compared to the Lacer predicted correct and error base quality profiles. For these, the target base quality was excluded; for example, an analysis of Q39 bases utilized only quality score histograms for Q6-Q38 and Q40. The fraction of single (unsupported) bases (out of the total number of nonconsensus, non-VCF bases; denoted f_Q;single_) was plotted for each quality score, and converted to a recalibration amendment as -10log_10_(f_Q;single_) for each quality score Q from 30-40.

### Quake analysis

Quake (version 0.3) (Kelley et al. 2010) was run on the FASTQ file for UTI89 using a k-mer of 13 and the --log option to output the bases and associated qualities that were corrected. Quake was similarly run on a downsampled (to 1%) FASTQ file of reads mapping to chr1 or chr19 of NA12878 using a k-mer of 15. The histogram of quality scores for bases corrected by Quake was taken as the error profile of bases identified by Quake.

### Program availability

Lacer is available at https://github.com/swainechen/lacer.

## Acknowledgments

This research was supported by the National Research Foundation, Prime Minister’s Office, Singapore under its NRF Research Fellowship Scheme (NRF Award No. NRF-RF2010-10) and the Genome Institute of Singapore (GIS)/Agency for Science, Technology and Research (A*STAR). We thank See Ting Leong, Dawn Choi and Xiaoan Ruan for the preparation and sequencing of the UTI89 Ion Torrent sequencing library. We thank Andreas Wilm, Niranjan Nagarajan, Shyam Prabhakar, and members of the Chen lab for helpful discussions and suggestions regarding the algorithm and manuscript. J.C.S.C. and S.L.C. designed the algorithm, performed the analyses, and wrote the paper.

## Supplemental Information

**Supplemental Figure S1: Lacer is insensitive to a variant database and an imperfect reference sequence.** (**A**) Quality score distributions for error bases in the UTI89 data set (using UTI89 as a reference) as predicted by Quake on the unmapped data (black) or by Lacer (red) and GATK (green) on the mapped data. When a perfect reference sequence is used (UTI89 resequencing data mapped to the UTI89 genome), the quality distribution for error bases is concordant among Lacer, GATK, and Quake. (**B**) Histogram of deviations of covariate quality scores for each data set indicated from the covariate quality scores derived from the GATK “gold standard” recalibration of the UTI89 data set using the UTI89 genome as the reference. For each covariate (context and cycle number), the difference in recalibrated quality between GATK (using the UTI89 genome as a reference) and a comparison recalibration was calculated. A histogram of all of these values is plotted here for GATK (using the MG1655 genome as a reference and w/ VCF; blue), Lacer (using the MG1655 genome as a reference; red), and Lacer (using the UTI89 genome as a reference; black) as the comparison recalibrations. Lacer is unaffected by changing the reference sequence, and the histogram of deviations is centered around zero, indicating no systematic bias in covariate recalibration. GATK using the MG1655 genome as a reference produces systematically lower recalibrated covariate quality scores, seen as a bias towards a tail of positive values. (**C**) Quality score distributions for error bases in the UTI89 data set (using MG1655 as a reference) as predicted by Quake on the unmapped data (black) or by Lacer (red) and GATK (w/ VCF; green) on the mapped data. Quake and Lacer produce concordant quality distributions for error bases. GATK produces a quality distribution that is biased towards high quality bases. (**D**) Lacer recalibration of the UTI89 data set using MG1655 (w/ or w/o VCF) as a reference. GATK recalibration of the UTI89 data set using UTI89 as a reference (w/o VCF) is shown for comparison. Provision of a variant database has minimal effect on the Lacer recalibration. (**E**) Lacer recalibration of the NA12878 chr1 data set (downsampled by region to 1Mbp) with and without excluding known variant positions. The provision of a database of known variants has little effect on Lacer recalibration. (**F**) Quality score distributions for error bases predicted by Quake on the unmapped NA12878 chr1 data set (black) or by Lacer (red) and GATK (green) recalibration on the mapped data. Quake and Lacer produce concordant quality distributions for error bases. GATK produces a quality distribution that is biased towards high quality bases. (**G**) Quality score distributions for error (blue) and correct (yellow) defined by GATK recalibration of the NA12878 chr1 data set. The quality score distribution for error bases is enriched for high quality scores and somewhat resembles the distribution for correct bases.

**Supplemental Figure S2: Validation of Lacer recalibration.** (**A**) Fraction of unsupported “single” non-reference, non-VCF bases with quality scores >=30. The fraction of single bases for UTI89 mapped to the UTI89 genome (circles) as a reference is ∼0.9. The fraction of single bases for the same UTI89 data set mapped to the MG1655 genome (triangles) is much lower, ∼0-0.2. The fraction of single bases for NA12878 chr1 mapped to GRCh37.p13 (squares) is also low, ∼0-0.1. (**B**) Histograms of coverage depth at all non-reference, non-VCF bases for the UTI89 (red) and NA12878 chr1 (green) data sets. (**C**) Correction of GATK recalibration using supporting base analysis for GATK recalibration on the UTI89 data set (UTI89 (left) and MG1655 (right) used as the reference sequence). Dark gray bars represent the difference between Lacer and GATK recalibration for each quality score. Light gray bars represent the quality difference from the GATK recalibration assuming only single (unsupported) non-reference, non-VCF bases are true errors. (**D**) Difference in VQS between the Lacer-recalibrated and uncalibrated call sets for all high quality (PASS; green) or low quality (LowQual; red) SNPs in the GATK-recalibrated call set for NA12878 chr1. On high quality SNPs, Lacer recalibration results in higher VQS (positive quality difference), while on low quality SNPs, the quality difference is centered about zero. (**E**) Difference in VQS between the GATK-recalibrated and uncalibrated call sets for all high quality (PASS; green) or low quality (LowQual; red) SNPs in the GATK-recalibrated call set. GATK recalibrated data results in generally lower VQS for all SNPs compared with uncalibrated data, but the decrease is greater for high quality SNPs than for low quality SNPs, contrary to expectation. (**F**) Distribution of overall base quality scores for the uncalibrated (black), Lacer-recalibrated (red), and GATK-recalibrated (green) NA12878 chr1 data. GATK recalibration results in a bimodal distribution that is not seen in the uncalibrated or Lacer-recalibrated data.

**Supplemental Figure S3: Recalibration of NA12878 chr19.** (**A**) Recalibration of the NA12878 chr19 sequencing data using GATK, with or without excluding known variants based on dbSNP, Mills and 1000 Genomes gold standard indels, and 1000 Genomes Phase 1 indels (w/ or w/o VCF, respectively), or using Lacer, w/o VCF or with dbSNP variant positions only (VCF positions only). GATK recalibration produces lower quality scores that are only partially increased by exclusion of known variants. Lacer recalibration using only known variant positions results in relatively similar recalibration to Lacer using the entire data set. (**B**) Distribution of quality scores for error bases predicted by Quake (black) on the unmapped data or by Lacer (red) and GATK (green) on the mapped data. Quake and Lacer produce concordant quality distributions for error bases. GATK produces a quality distribution that is biased towards high quality bases. (**C**) Distribution of overall base quality scores for the uncalibrated (black), Lacer-recalibrated (red), and GATK-recalibrated (green) data. None of the data sets have a bimodal distribution of base quality scores for chr19. (**D**) Venn diagram of Lacer- and GATK-recalibrated SNP calls identified by GATK HaplotypeCaller on chr19 that passed recalibration by the GATK VariantRecalibrator and that also matched the high-confidence NIST call set. Total numbers of SNP calls in each category and their associated Ti/Tv ratios are shown. SNP calls unique to Lacer recalibration have an aggregate Ti/Tv closer to the intersection than the SNP calls unique to GATK recalibration. (**E**) Comparison of VQS for SNPs identified in the uncalibrated, Lacer-recalibrated, or GATK-recalibrated data which matched the NIST call set. Lacer recalibrated data results in overall higher VQS than that from uncalibrated (red) and GATK recalibrated (green) data on these high confidence true positive SNPs. GATK recalibrated data results in overall lower VQS than uncalibrated (gray) data on these same SNPs. (**F**) Difference in VQS between the Lacer-recalibrated and GATK-recalibrated call sets for all high quality (PASS; green) or low quality (LowQual; red) SNPs in the GATK-recalibrated call set. Lacer recalibrated data results in a mild increase in VQS on low quality SNPs (as called by GATK recalibrated data), but this is small compared to the increase in VQS for high quality SNPs. Difference in VQS between the GATK-recalibrated and uncalibrated call sets for all high quality (PASS; green) or low quality (LowQual; red) SNPs in the GATK-recalibrated call set. GATK recalibrated data results in generally lower VQS for all SNPs compared with uncalibrated data, but the decrease is greater for high quality SNPs than for low quality SNPs, contrary to expectation. (**H**) Difference in VQS between the Lacer-recalibrated and uncalibrated call sets for all high quality (PASS; green) or low quality (LowQual; red) SNPs in the GATK-recalibrated call set. On high quality SNPs, Lacer recalibration results in higher VQS (positive quality difference), while on low quality SNPs, the quality difference is centered about zero.

**Supplemental Figure S4: Recalibration of non-human primate sequencing data.** Quality score distributions for error (red) and correct (green) defined by: (**A**) GATK recalibration (excluding known variants) or (**B**) Lacer recalibration of the Tibetan macaque chr1 sequencing data, and (**C**) GATK recalibration (excluding known variants) or (**D**) Lacer recalibration of the common marmoset exome sequencing data. For both data sets, the quality score distributions for error bases defined by GATK are enriched for high quality scores and somewhat resemble the distributions for correct bases. The quality profiles for Lacer were similar to those observed in previous data sets (c.f. **Fig. 1c**).

**Supplemental Figure S5: Recalibration of different NGS data sets using Lacer.** Empirical (recalibrated) quality scores plotted against reported (uncalibrated) quality scores for: (**A**) NA12878 chr1 exome (NCBI36 reference) sequencing data (Illumina GAIIx). Recalibrated quality scores are shown for each read group individually; (**B**) SRS017191 human metagenome (*Bacteriodes vulgatus* ATCC 8482 reference) sequencing data (Illumina GAIIx); (**C**) TruSeq Amplicon Cancer Panel (NCBI36 reference) targeted sequencing data (Illumina MiSeq); (**D**) UTI89 (UTI89 or MG1655 reference) sequencing data (Ion Torrent); (**E**) MG1655 (MG1655 or UTI89 reference) sequencing data (Roche 454). Recalibration by Lacer (w/o VCF) and GATK (w/ or w/o VCF) is shown. In the absence of a perfect reference, GATK recalibration produces lower quality scores compared with Lacer recalibration; these lower quality scores are only partially increased by exclusion of known variants. Furthermore, Lacer recalibration (using a perfect or imperfect reference) matches GATK’s recalibration when a perfect reference is provided. Lacer is therefore more accurate across a wide range of NGS data sets without the requirement of a variant database.

**Supplemental Table S1:** Impact of base quality score recalibration by Lacer and GATK on SNP calls from NA12878 chr19 (GATK HaplotypeCaller).

